# Intonation Units in spontaneous speech evoke a neural response

**DOI:** 10.1101/2023.01.26.525707

**Authors:** Maya Inbar, Shir Genzer, Anat Perry, Eitan Grossman, Ayelet N. Landau

## Abstract

Spontaneous speech is produced in chunks called Intonation Units (IUs). IUs are defined by a set of prosodic cues and occur in all human languages. Linguistic theory suggests that IUs pace the flow of information and serve as a window onto the dynamic focus of attention in speech processing. IUs provide a promising and hitherto unexplored theoretical framework for studying the neural mechanisms of communication, thanks to their universality and their consistent temporal structure across different grammatical and socio-cultural conditions. In this article, we identify a neural response unique to the boundary defined by the IU. We measured the EEG of participants who listened to different speakers recounting an emotional life event. We analyzed the speech stimuli linguistically, and modeled the EEG response at word offset using a GLM approach. We find that the EEG response to IU-final words differs from the response to IU-nonfinal words when acoustic boundary strength is held constant. To the best of our knowledge, this is the first time this is demonstrated in spontaneous speech under naturalistic listening conditions, and under a theoretical framework that connects the prosodic chunking of speech, on the one hand, with the flow of information during communication, on the other. Finally, we relate our findings to the body of research on rhythmic brain mechanism in speech processing by comparing the topographical distributions of neural speech tracking in model-predicted and empirical EEG. This qualitative comparison suggests that IU-related neural activity contributes to the previously characterized delta-band neural speech tracking.

## Introduction

Natural speech is prosodically structured: variations in volume, pitch, duration and other characteristics are an inherent part of utterances and in turn, of the linguistic signal that reaches our senses. Among its many roles, prosody underlies the sequencing of speech into speech chunks. Linguistic theory identifies a set of prosodic characteristics that fulfill the purpose of chunking speech cross-linguistically. These characteristics include an acceleration-deceleration dynamic of syllable delivery, resets in the slow modulations of pitch and volume, and at times – pauses (Chafe, 1994; Du Bois et al., 1992; Himmelmann et al., 2018; Seifart et al., 2021; Shattuck-Hufnagel & Turk, 1996). The resulting speech chunks are referred to as Intonation Units (henceforth IUs (Chafe, 1994; Du Bois et al., 1992); also Intonation(al) Phrases (Himmelmann et al., 2018; Seifart et al., 2021; Shattuck-Hufnagel & Turk, 1996), Intonation Groups (Cruttenden, 1997), Tone Groups (Halliday, 1967), and Elementary Discourse Units (Kibrik, 2019)). Since IUs are defined by a set of auditory cues, attuned listeners can identify them in languages that they do not understand (Chafe, 1994; Himmelmann et al., 2018). However, upon analyzing the progression of natural speech in context, an interesting meaning-related function of IUs emerges: IUs pace the flow of new information in the discourse in such a way that each IU appears to contain a maximum of one new idea. Supporting evidence for this comes from qualitative analyses in several languages (e.g., Chafe, 1987, 1994, 2018), and from discovering similar numbers of content items per IU across different languages (Himmelmann et al., 2018). An example transcript in Appendix S1 allows to appreciate the progression of IUs in a spontaneously recounted personal narrative (Labov & Waletzky, 1967). From an interactional linguistics perspective, IUs are also a useful resource for sequence organization in conversation and the construction of different speech actions (Bögels & Torreira, 2015; Ford & Thompson, 1996; Gravano & Hirschberg, 2011; Selting, 2010). Taking these properties together, IUs provide a window onto important functions of language cognition. In a recent study, we quantified the temporal structure of IU sequences in recordings of natural speech in six languages from around the world. We found that sequences of IUs form a ∼1 Hz rhythm in all these languages, despite substantial variation in their grammatical structures and socio-cultural profiles. The consistent temporal structure of IUs across languages provides further evidence for their importance in cognition (Inbar et al., 2020).

Research in cognitive neuroscience highlights an important role for temporal interactions between brain activity and speech. Neural activity tracks speech moment by moment at different time scales (Giraud & Poeppel, 2012; Gross et al., 2013). Temporal dynamics between different brain regions orchestrate this process according to attentional goals and demands (Mesgarani & Chang, 2012; Obleser & Kayser, 2019; Park et al., 2015; Zion Golumbic et al., 2013). Neural speech tracking has been found in the theta band (4-8 Hz). This temporal scale is thought to reflect neural tracking of syllables, which across languages tend to have a similar temporal structure: 4-8 syllables per second (Chandrasekaran et al., 2009; Ding et al., 2017; Greenberg et al., 2003; Pellegrino et al., 2011). Neural speech tracking at this rate promotes speech intelligibility (Ahissar et al., 2001; Ding & Simon, 2014; Peelle & Davis, 2012; Zoefel et al., 2018). Neural speech tracking has also been found in the delta band (1-2 Hz), however, there is less understanding as to what speech components are being tracked. Studies that investigate speech tracking of natural speech have associated delta band tracking with prosody (Gross et al., 2013; Keitel et al., 2017, 2018; Park et al., 2015). However, they do not present a detailed analysis of which prosodic modulations or what function they might serve in the language system. Another class of studies investigates tracking of synthesized language which is structured in time and purposefully lacks prosodic modulation (Bai et al., 2022; Ding et al., 2016). The finding in this case demonstrates convincing delta-band neural speech tracking of syntactic structure, in the absence of prosodic modulation. The interpretation of this finding raises two main difficulties. First, it is not obvious whether or how abstract linguistic structure in natural speech is structured in time, and given the substantial variation between language systems, it may be hard to generalize this type of finding to a cross-linguistic mechanism. Second, listeners perceive prosodic breaks at syntactic boundaries even when no acoustic cue for such a break exists (Buxó-Lugo & Watson, 2016; Cole et al., 2010). Thus, it is possible that delta-band neural speech tracking reflects some type of prosodic processing nonetheless (Gilbert et al., 2015; Glushko et al., 2022; Henke & Meyer, 2021).

We propose that IUs are powerful units with which to probe cognition during speech processing. They capture prosodic variation that is cross-linguistically relevant, and share temporal structure with previously found delta-band neural speech tracking. In order to enable a systematic study of the neural mechanisms of IUs, we use linguistic theory to identify these units in ongoing speech. Our analytic approach can be thought of as similar to the traditional event related potentials (ERPs) in EEG research. However, instead of experimentally triggering the brain to synthesized constructed speech we use the naturally occurring IUs as triggers, and investigate the neural response preceding and following this linguistic unit. In addition, we add an objective quantification of acoustic boundary strength at the word level to complement the subjective labeling of IUs. This measure of acoustic boundary strength enables us to characterize the IU impact on neural response while holding acoustic variation constant. Characterizing the neural response to IUs above and beyond acoustic boundary strength enables inference on the added cognitive and perceptual processes that are involved in the processing of speech.

Within the ERP literature, previous work using synthesized speech has identified the Closure Positive Shift (CPS) in response to prosodic phrase boundaries (Bögels et al., 2011; Steinhauer, 2003; Steinhauer et al., 1999). Prosodic boundaries in these studies rely on cues that are nearly identical to those defining IUs (cf. Chafe, 1994 to Steinhauer et al., 1999). The typical shape of this component is a centroparietal positivity in the EEG waveform at the offset of the last word in the phrase. Some studies also report a negative deflection in anterior electrodes, which at times begins even before the pause onset (Bögels et al., 2010; Pauker, 2013; Pauker et al., 2011). To date, the CPS has been documented in several languages from different phylogenetic units (‘families’) and geographical areas, including German, Dutch, English, Swedish, Japanese, Chinese, and Korean. Several studies investigated the prosodic, syntactic, and contextual conditions in which the CPS emerges. A pause in the speech signal is not necessary for the emergence of the CPS (Holzgrefe-Lang et al., 2016; Itzhak et al., 2010; Steinhauer et al., 1999), nor is any acoustic boundary cue whatsoever necessary if a boundary is predictable on the basis of syntactic structure alone (Itzhak et al., 2010). Correspondingly, a small CPS may emerge while reading written sentences, at the position of a comma (Steinhauer, 2003). In addition, responses to acoustically identical prosodic boundaries are modulated by the contextual predictability of the boundary (Kerkhofs et al., 2007). Taken together, these studies suggest that the CPS reflects a structuring of the input rather than a solely bottom-up response to the acoustic boundary cues giving rise to the prosodic phrasing. Nonetheless, a CPS also emerges in the absence of linguistic content at varying levels, suggesting a dependence on acoustics. A CPS may emerge at the closure of phrases that lack content words (jabberwocky sentences), semantics and syntax (pseudo-sentences), and even phonological content (hummed sentences) (Pannekamp et al., 2005). Pannekamp et al. (2005) showed that with less linguistic content, the CPS has a more anterior and right distribution. Another study showed that the magnitude of the CPS depends on the size of the boundary; the size of the boundary was manipulated by varying the duration of the final lengthening and the pause (Pauker, 2013). To the best of our knowledge, all studies concerning the CPS employ constructed and digitally manipulated sentences in isolated presentation conditions or with minimal context only. One exception is a recent study, that used reproduced speech segments that were originally recorded in natural environments (Anurova et al., 2022). However, this study included a pause-insertion manipulation at valid boundary positions as well as non-boundary positions. Manipulations such as these undoubtedly alter the listening mode in comparison to the natural experience of speech. By orienting the listener’s attention to prosodic breaks, the manipulation risks artificially raising the chances of finding an effect at prosodic boundaries (Bögels et al., 2011). Thus, the EEG response to prosodic boundaries in naturally occurring spontaneous speech is yet to be described.

In the present study, we model trial-level data using the Generalized Linear Model (GLM) framework in order to study the effect of IUs on the EEG response in natural listening conditions, on entirely spontaneous speech material (Figure 1). This modeling framework offers two key advantages over previous methods: (1) It treats the natural variation in acoustic boundary strength as a truly continuous variable (rather than collapsing over it to create a discrete set of experimental conditions), and (2) It allows us to ask whether EEG responses carry information on IUs above and beyond variations in acoustic boundary strength. We demonstrate (i) that IUs in natural Hebrew speech evoke an EEG response that is comparable in its timing and topographic distribution to the previously described CPS, and (ii) that this response is composed of two sub-components: one that is modulated by acoustic boundary strength and one that is unique to IU boundaries.

**Figure 1.**
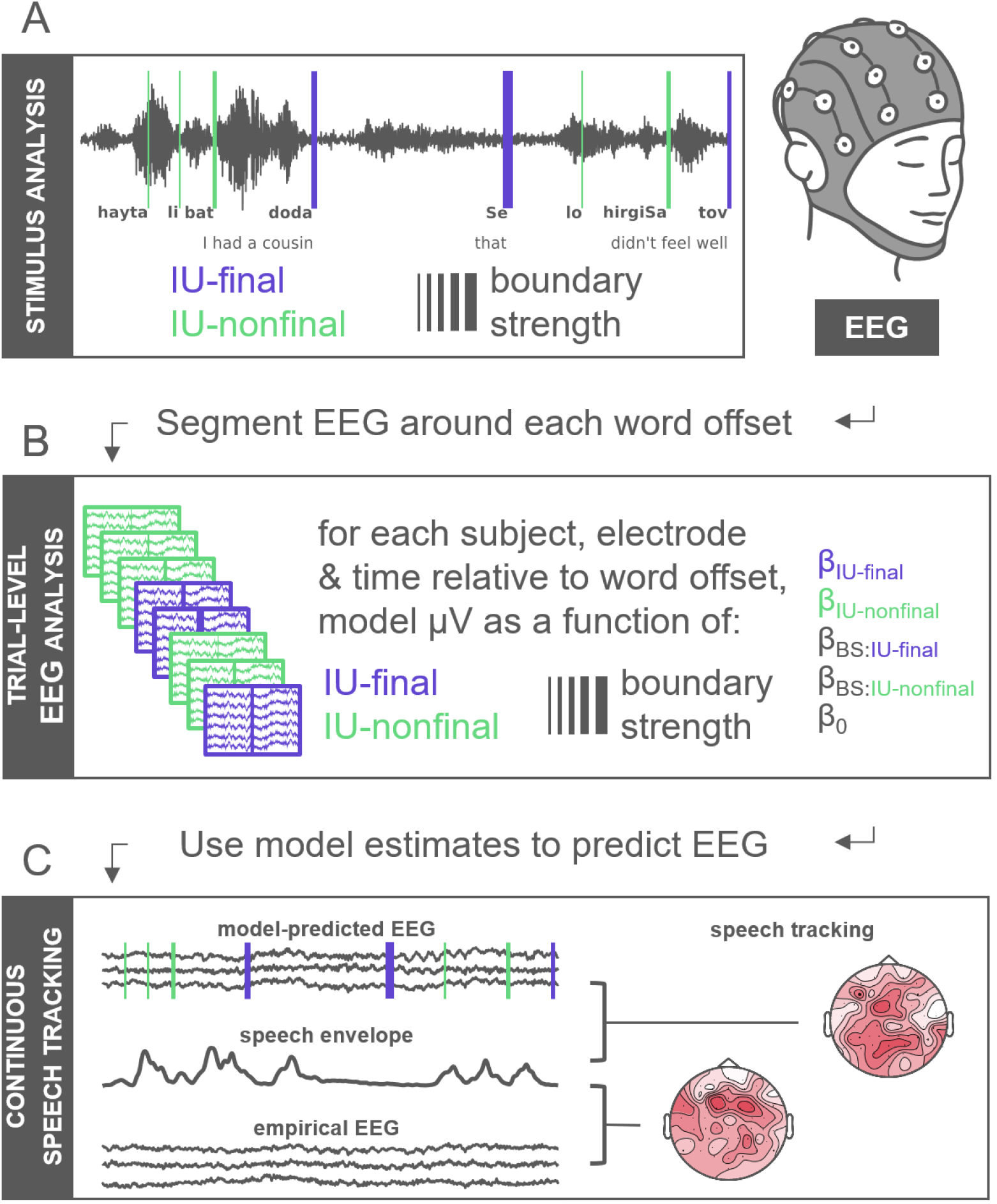
Methodology overview. (A) Spontaneous speech material in Hebrew was analyzed linguistically and acoustically (see Appendix S1 for an example transcript). Each orthographic word was timestamped (vertical lines relative to the audio waveform at word offsets), annotated as to whether it closed an IU or not (purple and green, respectively), and assigned an acoustic-based boundary strength score (illustrated by line width). Participants listened to this speech material while their EEG was recorded. (B) We segmented the EEG around word offset times, and modeled it as a function of IU closure and boundary strength (BS). (C) We use the model estimates and stimulus parameters to predict the continuous EEG response to the stimuli, and perform a descriptive neural speech tracking analysis on the model-predicted EEG and the empirical EEG relative to the speech envelope.

Based on our previous work demonstrating that sequences of IUs form ∼1 Hz rhythms (Inbar et al., 2020), we hypothesized that from a continuous perspective, the time course of responses to IUs would give rise to delta-band neural speech tracking. We use the estimated responses from the GLM model to predict the continuous EEG response to our stimuli and perform a neural speech tracking analysis on the empirical EEG and on the model-predicted EEG (Figure 1). This analysis reveals a similar topography of neural speech tracking in the delta band, but not in the theta band.

## Results

We analyzed EEG recordings of participants listening to different speakers describing an emotional life event. We transcribed the stories and analyzed them prosodically, into Intonation Units (IUs), and acoustically, computing a boundary strength score for each word based on a joint analysis of the speech envelope, fundamental frequency, and word duration information (see Materials and Methods). The distribution of boundary strength scores across two different word types, IU-final and IU-nonfinal, is presented in Figure 2A. As expected, words that close an IU tend to have stronger boundaries (β = 0.55, SE = 0.03, p < 0.001). For comparison, we also present the distribution of pauses across the different word types and the relation between pause duration and boundary strength score (Figure 2B and Figure 2C). It is noteworthy that pause duration accounts for at most 44.5% of the overall variance in boundary strength scores, and that 51.48% of IU-final words are followed by a pause of 50 ms or less (see Materials and Methods).

**Figure 2.**
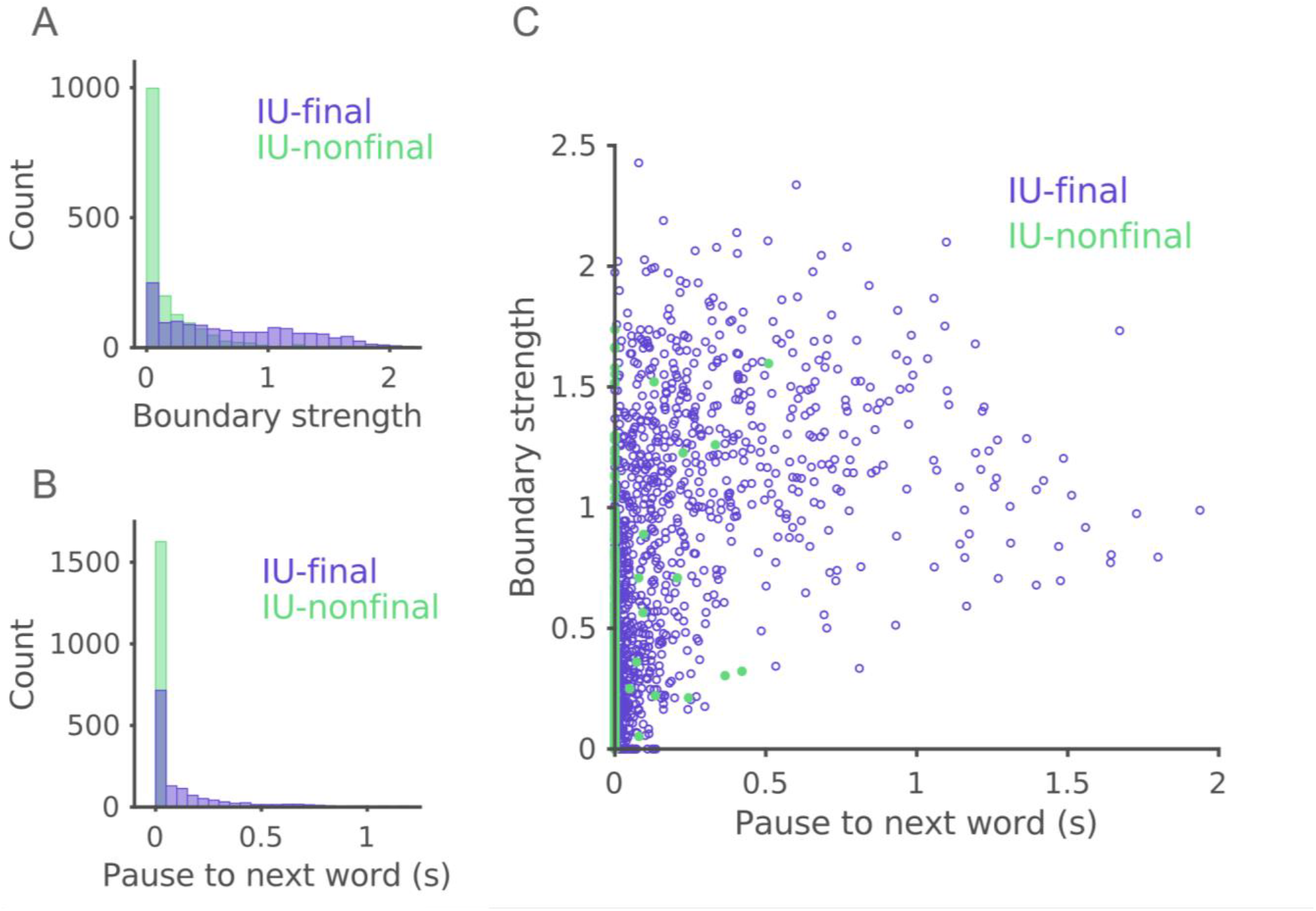
Relationship between boundary strength, pauses and IU closure. (A) Distribution of boundary strength scores in IU-final and IU-nonfinal words (purple and green, respectively). Histogram bins span 0.1 units of boundary strength. (B) Distribution of pauses from each word to the next in IU-final and IU-nonfinal words (purple and green, respectively). Histogram bins span 0.05 seconds. (C) Distribution of boundary strength scores relative to pause duration in IU-final and IU-nonfinal words (purple and green, respectively).

### EEG amplitude depends on IU closure and boundary strength

We modeled the contribution of two factors to the EEG response at a word offset: whether a word closed an IU or not (IU-final and IU-nonfinal, respectively), and its boundary strength. To this end, we used a hierarchical mass univariate general linear model (GLM) approach (Pernet et al., 2011; see Materials and Methods). In this framework, at the first level, the data of each participant is modelled using the IU closure predictor (a categorical predictor, IU-final or IU-nonfinal) and the boundary strength predictor (a continuous predictor, standardized across both levels of the categorical predictor). The effect of boundary strength is modeled separately in each level of the IU closure condition to accommodate for the possibility that the effect differs between IU-final and IU-nonfinal words (see Materials and Methods). Beta coefficients are estimated for each electrode and time point. At the second level, beta estimates are entered into a non-parametric cluster-based analysis to evaluate the significance of these effects at the group level over time and scalp location.

This analysis revealed an effect of IU closure and an effect of boundary strength in IU-final words on the EEG response around word offset. First, for words with equivalent and average boundary strength, words that close an IU are associated with a centroparietal positivity preceded by a right-anterior negativity compared with words that do not close an IU. This was supported by the significant effect of IU closure (Figures 3A-D; negative cluster extending from -76 to 225 ms following word offset, p = 0.009; positive cluster extending from 133 to 504 ms following word offset, p < 0.001). There was an additional positive cluster preceding the right-anterior negativity, circa 500 ms before word offset (Figure 3A; cluster extending from -545 until -449 ms relative to word offset, p = 0.006). It is noteworthy that the IU closure contrast provided equivalent results when performed on non-standardized variables (Figure S1A-D). In essence, this means that the effect of IU closure is also apparent when comparing IU-final and IU-nonfinal words with zero boundary strength rather than average boundary strength. Second, for words that close an IU, stronger boundaries elicited a larger anterior negativity (Figures 3E-G; post hoc test for boundary strength effect, two clusters extending from 119 to 311 ms and from 324 to 400 ms following word offset, p < 0.001 and p = 0.004, respectively). Here too there was a positive cluster preceding the negativity, circa 200 ms before word offset (Figure 3E; cluster extending from -197 until -141 ms relative to word offset, p = 0.006). For words that do not close an IU, the effect of boundary strength was only trending (Figure S1E-G), perhaps due to the overall skewed distribution of boundary strength scores within IU-nonfinal words (Figure 2A). Finally, we tested whether the effect of boundary strength differed between IU-final and IU-nonfinal words and found no significant difference (see Materials and Methods).

**Figure 3.**
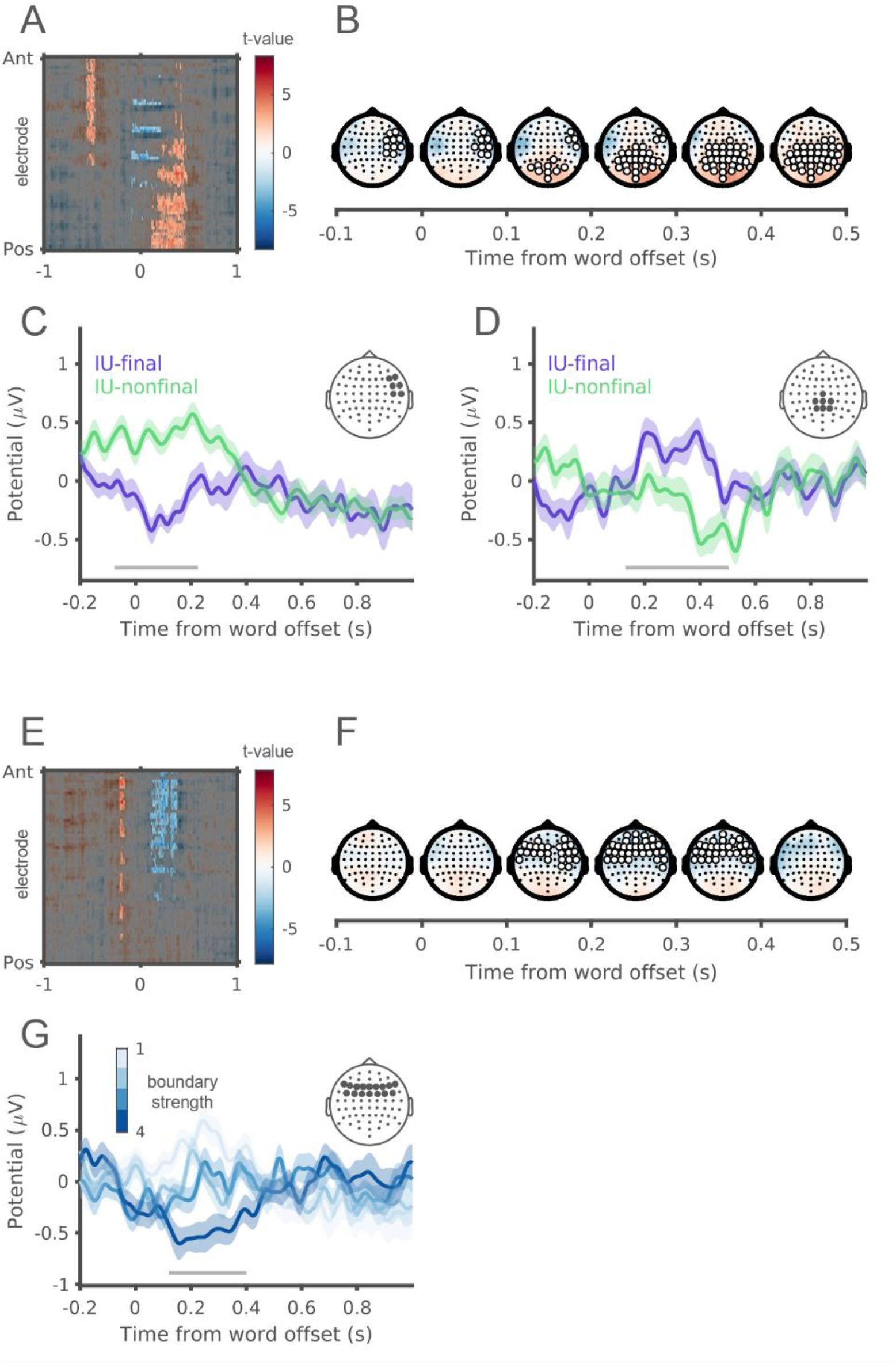
EEG amplitude depends on IU closure and boundary strength. (A) The full spatiotemporal results for the IU closure effect. A t-value is calculated for the observed difference between IU-final and IU-nonfinal EEG responses at each electrode and timepoint in a 2-second window around word offset. Cluster-based bootstrap tests reveal significant clusters at the group level (unmasked). (B) Topographical distributions of the IU-closure contrast t-values, from -100 ms to 500 ms around word offset in 100 ms steps. Electrodes in significant clusters are highlighted in white if they span 15 consecutive milliseconds. (C) ERP traces in response to IU-final and IU-nonfinal words with mean boundary strength score (purple and green, respectively), illustrating the right-anterior-negative cluster. The traces show the grand average over the EEG electrodes highlighted in the inset topography. Shaded ribbons correspond to ±1 SEM. The horizontal gray bar marks the timepoints over which this cluster was significant in the GLM model. (D) ERP traces in response to IU-final and IU-nonfinal words with mean boundary strength score (purple and green, respectively), illustrating the centroparietal positive cluster. The traces show the grand average over the EEG electrodes highlighted in the inset topography. Shaded ribbons correspond to ±1 SEM. The horizontal gray bar marks the timepoints over which this cluster was significant in the GLM model. (E) The full spatiotemporal results for the effect of acoustic boundary strength within IU-final words. A t-value is calculated for the estimated change in EEG response against zero at each electrode and timepoint in a 2-second window around word offset. Cluster-based bootstrap tests reveal significant clusters at the group level (unmasked). (F) Topographical distributions of the effect of boundary strength within IU-final words, from -100 ms to 500 ms around word offset in 100 ms steps. Electrodes in significant clusters are highlighted in white if they span 15 consecutive milliseconds. (G) ERP traces in response to IU-final words with different levels of boundary strength, illustrating the anterior negative cluster. Four different levels are presented, corresponding to quartiles of boundary strength scores within IU-final words. The traces show the grand average over the EEG electrodes highlighted in the inset topography. Shaded ribbons correspond to ±1 SEM. The horizontal gray bar marks the timepoints over which this cluster was significant in the GLM model.

### Modeling ongoing EEG response for the computation of speech tracking

In a previous study, we characterized the temporal structure of IU sequences and showed that they form a ∼1 Hz rhythm (Inbar et al., 2020). From a continuous perspective, the time course of responses to IUs should therefore give rise to neural activity at that temporal scale. Previous studies investigating ongoing speech have, in fact, demonstrated neural tracking of speech within the delta band (< 2 Hz). In order to link our findings to the neural tracking literature we predict (i.e., reconstruct) the ongoing neural response to the stories from the model estimates. Using an information theoretic framework (Ince et al., 2017), we quantify the relation of the continuous EEG to the speech envelope in two bands, delta (0.8-1.1 Hz) and theta (3.5-5 Hz; Figure 4A). First, we find that the group-average mutual information (MI) between the model-predicted EEG responses and the speech envelope was significantly higher than the MI of the empirical EEG response and the speech envelope. In both delta and theta, a two-tailed two-sample Kolmogorov-Smirnov test rejected the hypothesis that the MI of the speech envelope with the model-predicted EEG and the MI of the speech envelope with the empirical EEG are from the same continuous distributions (delta band: *D* = 1, p < 0.001; theta band: *D* = 1, p < 0.001). To compare the topographical distributions of MI despite this difference in magnitude we transformed MI values in each topography into z-scores. Model-predicted EEG responses to IUs from our model capture similar-looking relationships to the speech envelope in the delta band, but not in the theta band (Figure 4B). This observation suggests that previous findings on delta-band neural speech tracking are related to the processing of Intonation Units.

**Figure 4.**
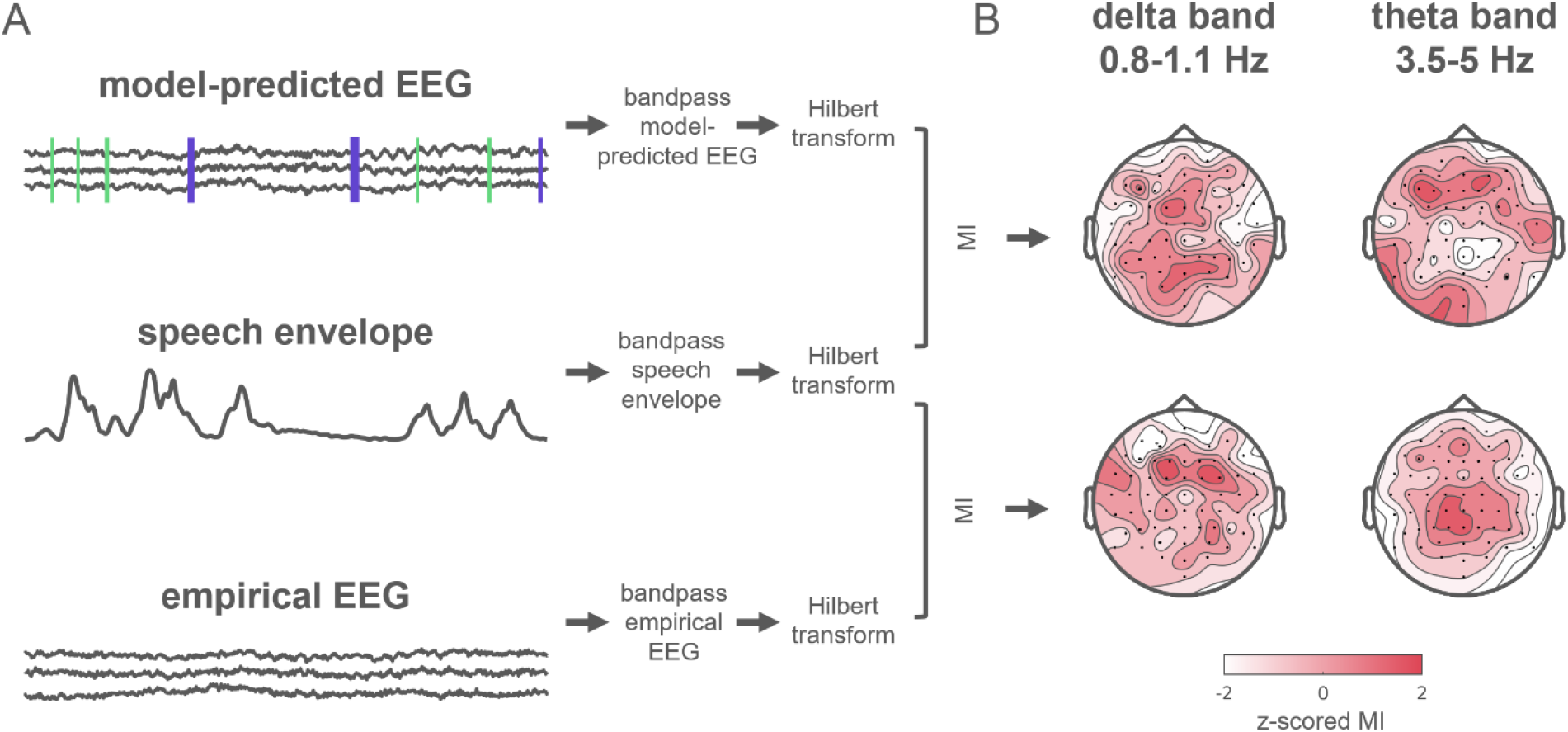
Investigating neural speech tracking using model-predicted EEG. (A) Neural speech tracking pipeline, quantifying mutual information (MI) between model-predicted EEG and the speech envelope, as well as between the empirical EEG and the speech envelope. (B) Z-scored MI values of the model-predicted EEG with the speech envelope (top) and the empirical EEG with the speech envelope (bottom), quantifying shared information in the delta band (left) and theta band (right). MI values were z-scored within each topography.

## Discussion

We set out to study the neural response to Intonation Units (IUs) with EEG. IUs are prosodically-defined units that serve as a window onto important functions of the language system. They pace the information in speech and serve as a resource for organizing conversational sequences. They are found across all human languages and have a common temporal structure. The results of the current study suggest that the neural system is attuned to these units, a finding that opens new paths for investigating the neural substrates of language from a usage-based linguistics perspective.

The EEG response at IU closure includes a negative deflection at right-anterior electrodes, starting as soon as the last word in the IU ends and lasting circa 200 ms. IU closure is further characterized by a centroparietal positive deflection between 150-500 ms after the last word in the IU. Within words that close an IU we found that stronger acoustic boundaries elicit a larger anterior negativity between 100-400 ms. This was also the direction of the trend in words that do not close an IU (Figure S1E-G).

The neural response to IUs tightly corresponds to the CPS component previously described for phrase boundaries. Our findings add to this literature in several ways. First, we describe neural responses while participants listened to speech in naturalistic conditions, in contrast to the isolated, constructed sentences that are typically used as stimuli in the CPS literature. Second, our work identifies what appear to be two different components within the classical CPS: a late anterior negativity that is dependent on boundary strength, and a centroparietal positivity and early- and right-anterior negativity that are not dependent on boundary strength. In this regard, we obtained results that differ from those of Pauker (2013), in which both anterior negativity and centroparietal positivity did depend on boundary strength. The difference might stem from any of different experimental procedures, including the different language of presentation, radically different stimulus types, and a different operationalization of boundary strength. Here we relied on rich natural speech material and operationalized boundary strength using a measure that includes prosodic information beyond that considered by Pauker (i.e., pitch modulation). Note that Pauker suggested that further study of the negativity is required, and the results of the current study bear out this suggestion and point to what appear to be two sub-components within the CPS.

IU boundaries are defined auditorily, and hence depend (albeit subjectively) on acoustic properties. On the other hand, in naturally occurring spontaneous speech IUs assume a role of organizing the speech stream in units – pacing ideas and pacing conversational actions. We attempt to address this complex manifestation of prosodic phrase boundaries in naturally occurring spontaneous speech. First, we implement an algorithm for measuring acoustic-based boundary strength as a continuous variable. Next, we quantify the relation of boundary strength to expert transcription of natural speech to IUs. Finally, while previous studies that characterized EEG at prosodic phrase boundaries tackle the acoustic and organizational facets separately, with our modelling approach we do so in tandem. Our approach and results suggest that to the extent that listeners are capable of listening to speech in a foreign language attentively enough, non-speakers of a language would show a boundary strength effect yet not an IU closure effect.

Both IU closure and stronger acoustic boundaries within words that close an IU were associated with a positive deflection that preceded word offset by hundreds of milliseconds (circa 500 ms in the former, and 200 ms in the latter). Especially in the case of the IU closure contrast such an early response might pertain to processes related to word onset, as the duration of words in our stimuli was 373±214 ms (M±SD). For example, there is evidence that preceding and following word onset, listeners’ brains are engaged in next-word prediction and surprise (or lack-of-surprise) at the incoming word (e.g., Goldstein et al., 2022). The current study focused on responses to word offsets in accord with the mainstream CPS literature, but future studies may investigate this putative word onset response and clarify whether there is, for example, a different engagement in word prediction depending on whether a word closes an IU or not.

In a final step, we related our results from the GLM model to the body of research on rhythmic brain mechanisms in speech processing. We used the estimated responses from our model to predict continuous EEG responses, and subsequently computed mutual information with the speech envelope in this signal and in the empirical EEG. Comparing the topographical distributions of neural speech tracking in the model-predicted and empirical EEG, we find similar topographies of neural speech tracking in the delta band but less so in the theta band. This observation, though qualitative, suggests that IU-related neural activity contributes to the previously characterized delta-band neural speech tracking. We note that a link between delta-band neural speech tracking and the CPS has been previously conceptualized (Meyer et al., 2016, 2020) but to the best of our knowledge has never been investigated. Therefore, in this final step, the current study also bridges two fruitful perspectives on language processing: the single evoked-response and the continuous process. We demonstrate that characterizing evoked responses locked to naturally occurring events can equip the analysis of continuous speech with powerful theoretical models. This approach can readily advance the study of neural mechanism of ongoing cognition in different domains by incorporating explicitly modelled events. Within the domain of speech, the focus on IUs embraces the natural co-variation of acoustic boundary strength and unit-ness. By doing so we promote a view of the language system as an embodied system whereby abstract structure supervenes upon perceptual modulations in time (Kreiner & Eviatar, 2014).

## Materials and methods

We analyzed EEG recordings of participants listening to different speakers describing an emotional life event. This data was collected as part of an independent project and summarized in detail in Genzer, Ong, Zaki and Perry (2022).

### Participants

Genzer and colleagues (Genzer et al., 2022) recorded EEG from 57 Hebrew-speaking undergraduate students from the Hebrew University of Jerusalem. The participants received monetary compensation at a rate of 40 NIS per hour (∼$15) or course credit. All participants reported normal or corrected to normal visual acuity and had no history of psychiatric or neurological disorders. Participants gave their informed consent before the experimental session. The study was approved by the institutional review board of ethical conduct at the Hebrew University of Jerusalem.

### Stimuli

Each participant listened, through speakers, to three out of nine stories from an Israeli Empathic Accuracy stimulus set (Jospe et al., 2020). In this stimulus set, Hebrew speakers described emotional life events as they sat in front of a professional recorder. The duration of the stories was between 2:01 and 3:48 minutes, with an average duration of 2:43 minutes. The nine stories were grouped into sets of three such that the sets were of approximately equal duration (range: 454–520 s). The assignment of story-set to participant was random and counterbalanced.

Details on the *Experimental design* and *EEG acquisition* can be found in Genzer et al. (2022).

### Stimulus annotation

Two trained native Hebrew speaking annotators transcribed the 9 stories that served as stimuli, and segmented them into IUs according to the criteria devised by Chafe and colleagues (Chafe, 1994; Du Bois et al., 1992). The segmentation process involves close listening to the rhythmic and melodic cues for IU boundaries, as well as performing acoustic analyses in PRAAT (Boersma & Weenink, 2022) for the extraction of pitch contours which are used to support perceived resets in pitch. We previously described this process in Inbar et al. (2020). The annotators checked each other’s segmentations and reached consensus at ambiguous points.

Next, we time-stamped each orthographic word in the transcription relative to the recording, and coded whether it appeared at the end of an IU or not, according to the segmentation described above. The timestamping was done in PRAAT (Boersma & Weenink, 2022), imported into ELAN (2022), and exported from there into a single text file including all the words from all recordings. We processed this file with the aid of custom-written scripts in R (R Core Team, 2022), and created a table including word onset and offset times, word duration, pause duration from each word to the next, and a boundary strength score (see next section: Boundary strength score). Table 1 presents summary information about the recordings and the counts of words and IUs in each recorded story.

**Table 1.**
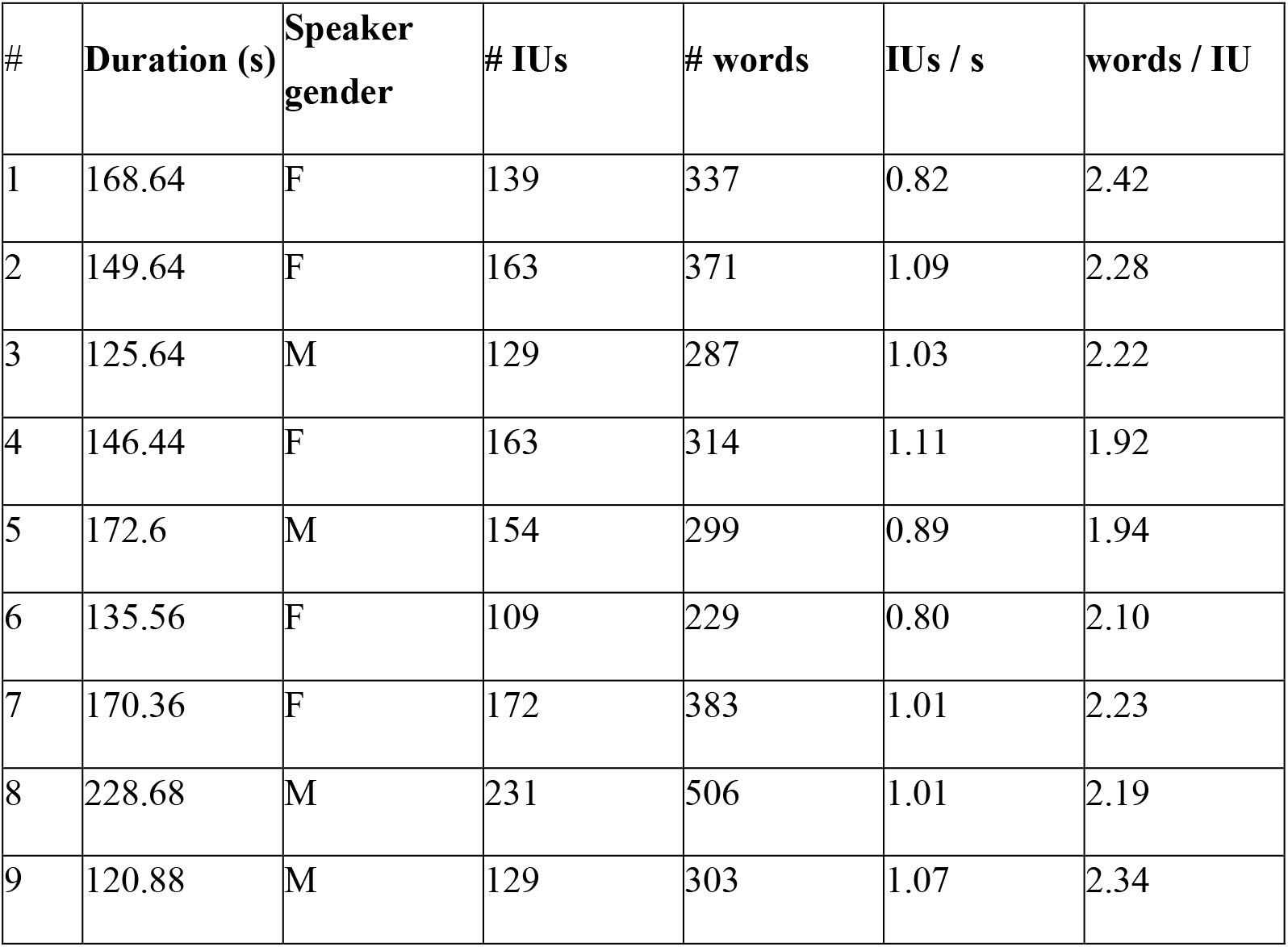
Summary information of the speech recordings and annotations.

| # | Duration (s) | Speaker<br>gender | # IUs | # words | IUs / s | words / IU |
| --- | --- | --- | --- | --- | --- | --- |
| 1 | 168.64 | F | 139 | 337 | 0.82 | 2.42 |
| 2 | 149.64 | F | 163 | 371 | 1.09 | 2.28 |
| 3 | 125.64 | M | 129 | 287 | 1.03 | 2.22 |
| 4 | 146.44 | F | 163 | 314 | 1.11 | 1.92 |
| 5 | 172.6 | M | 154 | 299 | 0.89 | 1.94 |
| 6 | 135.56 | F | 109 | 229 | 0.80 | 2.10 |
| 7 | 170.36 | F | 172 | 383 | 1.01 | 2.23 |
| 8 | 228.68 | M | 231 | 506 | 1.01 | 2.19 |
| 9 | 120.88 | M | 129 | 303 | 1.07 | 2.34 |

### Boundary strength score

Each orthographic word was given a score that indicated the strength of the boundary at its offset. The score was estimated using a published algorithm – the Wavelet Prosody Toolkit (Suni et al., 2017). This algorithm derives fundamental frequency and intensity information from the speech audio signal, and duration information from time-stamped word annotations. Thus, the algorithm operates precisely on the signals that correspond to the main cues for an IU boundary. These signals are summed to one and decomposed to several scales using Continuous Wavelet Transform. The algorithm then finds peaks and troughs in the output of the Continuous Wavelet Transform. Based on these peaks and troughs, the algorithm defines for each annotated word (see above) a prominence value and a boundary value (of which we only used the boundary value). In general, the prominence value of a word is defined to be the strongest peak within the word. The boundary value of a word is defined to be the strongest trough between two peaks: the strongest peak within the word and the strongest peak within the next word.

We provided as input to the algorithm the speech audio files and the time-stamped word annotations, and applied the algorithm with default settings. Our procedure deviated from that described in Suni et al. (2017) in a single respect: since we annotated breaths in the stories as single-word IUs, in our application of the algorithm we also assigned boundary scores to breaths.

### Relation between IU closure, boundary strength scores, and pause durations

To quantify the difference in boundary strength scores between IU-final words and IU-nonfinal words we fitted a mixed-effects linear model, predicting boundary strength score from IU closure condition. The model included a by-story random intercept as well as a by-story random slope for the effect of IU closure condition (IU-final and IU-nonfinal), to account for slight variations between the stories. The model’s explanatory power related to the fixed effect alone is 0.30. The model’s intercept, corresponding to the average boundary strength score of IU-nonfinal words, is estimated at 0.16, 95% CI [0.13, 0.18]. IU-final words have, on average, a significantly larger boundary strength score, with a difference estimated at 0.55 [0.49, 0.61], t(3023) = 17.07, p < .001).

To quantify the relationship between boundary strength scores and pause durations we fitted a mixed-effects linear model, predicting the standardized boundary strength scores from the standardized durations of the pauses from each word to the next. The model included a by-story random intercept as well as a by-story random slope for the effect of pause duration, to account for slight variations between the stories. The model’s total explanatory power is 0.45 (conditional R^2) and the part related to the fixed effect alone (marginal R^2) is 0.36. The model’s intercept, corresponding to the average boundary strength score for the mean pause duration (mean and not zero because the variables were standardized), is estimated at 0.01 [-0.04, 0.07]. The pause to the next word is significantly and positively related to the boundary strength score (beta = 0.67, [0.46, 0.88], t(3014) = 6.25, p < .001).

Mixed-effect models were estimated using REML and `nloptwrap` optimizer with `lme4` (Bates et al., 2015) in R (R Core Team, 2022). Conditional and marginal R^2 were calculated with the aid of the `MuMIn` package (Barton, 2022). 95% CIs and p-values were computed using a Wald t-distribution approximation with the aid of the `report` package (Makowski et al., 2021).

### EEG preprocessing

We processed the 64-channel EEG in MATLAB (The MathWorks, R2018b) using the FieldTrip toolbox (Oostenveld et al., 2011) and custom scripts. We re-referenced the signal in the EEG channels to the average signal from the mastoid electrodes. We computed bipolar derivations of the horizontal and vertical pairs of electrodes around the eyes and appended the resulting horizontal and vertical EOG signals to the EEG channels. We segmented the data to blocks according to the presented clips, starting from two seconds before the story began until two seconds after the story ended. We removed slow drifts in the EEG using spline interpolation as implemented in the function `msbackadj` (part of the MATLAB Bioinformatics toolbox) and similarly to (Ofir & Landau, 2022). The window size was 4 seconds, with a step of 0.75 second between two consecutive windows. The median of each window was taken as the baseline value. We detected large artifacts in the EEG semi-automatically with the aid of the function `ft_artifact_zvalue`, padding each artifact with 500 msec on either side. After visually inspecting the detected artifacts, we substituted them with zeros. We detected and removed excessively bad channels by visual inspection. We then ran ICA using the infomax algorithm (maximum 1024 training steps, following PCA, and saving 20 components). We identified components corresponding to eye- and muscle-related activity and removed them from the data.

We replaced rejected electrodes with the plain average of all neighbors as implemented in `ft_channelrepair`. Next, we defined trials as epochs from 1000 msec before to 1000 msec after each word offset. Trials that overlapped with the semi-automatically detected artifacts were discarded. We removed the mean from each epoch as a baseline. We removed remaining artifactual trials after visual inspection guided by the summary statistics in the `ft_rejectvisual` function. In total, we excluded 7 participants from further analyses: the data of 5 participants included recording failures (and was not submitted to preprocessing in the first place), and the data of 2 participants included excessive amounts of artifacts that affected >15% of their trials. In the final sample of 50 participants, the average number of rejected ICA components was 2.74 ± 1.32 (M ± SD), the average number of interpolated channels was 1.1 ± 1.47 (M ± SD; Max: 6), and the average number of remaining trials was 966.5 ± 72.72 (M ± SD; Range: 854-1072).

For visualizing the effect of IU closure (IU-final vs. IU-nonfinal) for mean boundary strength (Figures 3C and 3D), the time-domain data was processed as follows. After we removed artifactual trials, we standardized the per-trial boundary strength scores of each subject. Next, for each IU closure condition we selected trials with a boundary strength z-score between ± 0.5. This resulted in 138.46 ± 11.30 and 138.7 ± 9.9 (M ± SD) trials per participant per IU closure condition, (IU-final and IU-nonfinal, respectively; a two-tails paired-samples t-test indicated no significant difference between the trial counts, t(49) = -0.168, p = 0.87). We averaged the data in each trial over two subsets of electrodes (6 electrodes for the negative cluster: F8, FT8, T8, F6, FC6, C6; 7 electrodes for the positive cluster: Cz, CPz, Pz, CP1, CP2, P1, P2), based on previous literature and in accord with the significant clusters we found in the statistical modelling. Next, we applied an FIR windowed-sinc zero-phase lowpass filter, with 20 Hz as the cut-off frequency. Finally, we averaged the trials within IU closure condition and within-participant, and then across participants. For visualizing the effect of IU closure (IU-final vs. IU-nonfinal) for zero boundary strength (Figures S1C and S1D), the time-domain data was processed identically except the trial selection, which was done as follows. We binned the entire sample of boundary strength scores into 8 equally populated bins (quantiles). For each participant, we selected those trials with a boundary strength score in the first quantile, corresponding to those trials with the weakest boundary strength score. This resulted in 52.64 ± 9.3 and 229.76 ± 21.07 (M ± SD) trials per participant per IU closure condition, (IU-final and IU-nonfinal, respectively; a two-tails paired-samples t-test indicated a significant difference between the trial counts, t(49) = -55.6063, p < 0.0001).

For visualizing the effect of boundary strength within IU-final words (Figure 3G), the time-domain data was processed as follows. First, for IU-final words only, we binned the entire sample of boundary strength scores into quartiles. We averaged the data in each IU-final trial over a subset of electrodes (18 electrodes: Fz, F1, F2, F3, F4, F5, F6, F7, F8, FCz, FC1, FC2, FC3, FC4, FC5, FC6, FC7, FC8), based on the significant cluster we found in the statistical modelling. Next, we applied an FIR windowed-sinc zero-phase lowpass filter, with 20 Hz as the cut-off frequency. Finally, we averaged the trials within quartile and within-participant, and then across participants. For visualizing the effect of boundary strength within IU-nonfinal words (Figure S1G), the time-domain data was processed as follows. First, for IU-nonfinal words only, we binned the entire sample of boundary strength scores into 8 quantiles. Since many IU-nonfinal words have the same zero value and therefore cannot be placed in separate bins, the operation resulted in only 6 quantiles. Roughly applying a median split, we separately grouped the three-lower and three-upper quantiles of boundary strength scores. We averaged the data in each IU-nonfinal trial over a subset of electrodes (18 electrodes: Fz, F1, F2, F3, F4, F5, F6, F7, F8, FCz, FC1, FC2, FC3, FC4, FC5, FC6, FC7, FC8), based on the significant cluster we found in the statistical modelling of the effect of boundary strength score in IU-final words (Figure 3G). Next, we applied an FIR windowed-sinc zero-phase lowpass filter, with 20 Hz as the cut-off frequency. Finally, we averaged the trials within split and within-participant, and then across participants.

### Statistical modeling

We analyzed the 2-second windows around each word offset using the hierarchical mass univariate general linear modelling approach implemented in LIMO (Pernet et al., 2011). At the first level, we modeled the EEG amplitude at each electrode and timepoint combination per participant using the following formula:

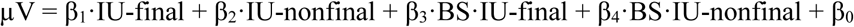

This model includes a predictor for IU closure condition (IU-final vs. IU-nonfinal) and a continuous, linear predictor for boundary strength (BS), with a different slope depending on IU closure condition. The predictor IU-final was 1 for all words that closed an IU, and 0 otherwise. The predictor IU-nonfinal was 1 for all words that did not close an IU, and 0 otherwise. Boundary strength scores were standardized across both IU closure conditions per participant, such that boundary strength 0 is the mean boundary strength across all words. The model parameters, then, have the following interpretation:

1. β^1^ – the effect of a word with mean boundary strength that closes an IU.
2. β^2^ – the effect of a word with mean boundary strength that does not close an IU.
3. β^3^ – the effect of increasing the boundary strength by 1 standard deviation for IU-final words.
4. β^4^ – the effect of increasing the boundary strength by 1 standard deviation for IU-nonfinal words.
5. β^0^ – the predicted EEG amplitude after regressing out all other effects.

In total, 64 X 1,025 models (65,600, one for each electrode and timepoint combination) were fitted to the data of each participant. The model parameters were estimated using a weighted least squares method for each participant separately. After the model parameters were estimated for each timepoint and electrode for each participant, we computed two contrasts, that is, linear combinations of parameters that capture comparisons of interest, henceforth, effects. The main contrast of interest was the IU closure effect, which we computed as the difference β^1^ – β^2^ (responses to IU-final words minus responses to IU-nonfinal words, when the boundary strength equals the mean for that dataset). We computed an additional contrast, the difference β^3^ – β^4,^ to capture any difference between the slope of the regression against boundary strength for IU-final vs. IU-nonfinal words. Finally, we tested the effect of boundary strength separately for IU-final (β^3^) and IU-nonfinal (β^4^) words.

As the 4 effects were computed at each electrode and timepoint for each participant, a single participant is represented by 4 spatiotemporal matrices of size 64 X 1,025, one per effect. Every element in each matrix, that is, every electrode-timepoint pair of a participant, is a single observation in the statistical significance test of a specific effect. The significance of the contrasts is estimated at the group level, using a clustering approach to correct for multiple comparisons and following LIMO conventions. For the two contrasts (IU closure effect and the difference in boundary strength slopes between the different levels of the IU closure condition) we used an alpha of 0.05. For the effects of boundary strength within IU-final and IU-nonfinal trials we used an alpha of 0.025, to correct for post-hoc comparisons.

### EEG prediction from GLM estimates

We used the GLM parameter estimates and the stimulus annotations to predict the continuous EEG signal in response to the stimuli. To this end, we extracted for each participant their estimated model parameters (which include β^1^, β^2^, β^3^, β^4^ and β^0^ parameters for each electrode and timepoint combination) and their predictor matrix, which includes the IU closure annotation and boundary strength score of each word that the participant heard as part of the recordings. By matrix-multiplying the estimated parameter matrices and the predictor matrix we obtained 2-second predicted EEG responses, centered on the offset of each word. We then added the windows consecutively to a zero-filled EEG time course according to the original offset times of each word in the recording. Note that the model-predicted EEG was calculated using the model estimates of each participant and the specific stimuli they were presented. Therefore, 50 distinct full EEG datasets were constructed, one for each participant.

### Neural speech tracking computation

We perform a neural speech tracking analysis in the original EEG recordings and in the predicted EEG that we constructed based on the GLM model estimates and the stimulus parameters. To this end, we followed common procedures in the field, including computing the speech envelope, focusing on two bands of interest, and using the information theoretic measure of Mutual Information (MI) to operationalize neural speech tracking (Ince et al., 2017).

The speech envelope is a representation that captures amplitude fluctuations in speech. It has been used in many studies as the signal to which ongoing neural fluctuations are compared to establish neural speech tracking. To compute it, we follow the steps presented in Gross et al. (2013) following Chandrasekaran et al. (2009). Specifically, each recording was converted from stereo to mono by averaging the two channels, and subsequently downsampled to 20kHz. Each recording was band-pass filtered into 10 bands between 100-10,000 Hz, with cut-off points designed to be equidistant on the human cochlear map. Amplitude envelopes were computed for each narrow band as absolute values of the Hilbert transform. These narrowband envelopes were downsampled to 1,000 Hz and subsequently averaged, yielding the wideband envelope. The wideband envelope was smoothed using a 50 ms sliding Gaussian filter and divided by its maximal value to be on a scale of 0-1.

Next, we perform a neural speech tracking analysis in two frequency bands of interest: the delta band, between 0.8-1.1 Hz, and the theta band, between 3.5-5 Hz. These cutoff frequencies conform with recent neural speech tracking literature (e.g., Kaufeld et al., 2020), and in the case of the delta band, correspond to the actual minimal and maximal IU rate in our stimuli (see Table 1). We filtered the speech envelopes, the empirical EEG and the predicted EEG in the two frequency bands using forward and reverse 2^nd^ order butterworth filters. Then, for each bandpassed signal (speech envelope, empirical EEG and model-predicted EEG in both delta band and theta band), we computed the complex representation of the Hilbert transform to obtain instantaneous information about both phase and amplitude. We calculated Mutual Information using the Gaussian copula approach (Ince et al., 2017) to quantify the relationship between the instantaneous phase and amplitude of the speech envelope in a single band and the instantaneous phase and amplitude of the time-delayed EEG signals in the same band (original and reconstructed). Conforming with previous literature (e.g., Bröhl & Kayser, 2021; Kaufeld et al., 2020; Keitel et al., 2018; Park et al., 2015), we quantified the relationship using 5 lags between the stimulus and the EEG, ranging from 60 to 140 ms in steps of 20 ms, and subsequently averaged the MI over the 5 lags. We performed this MI computation in the two bands for each of the three stories each participant listened to, and subsequently averaged the MI across the stories within each participant, within band. This was done separately for the MI with the empirical EEG and the MI with the predicted EEG. Finally, we computed the grand average MI across participants for the empirical EEG and the predicted EEG, separately for the delta band and the theta band.

## Code and data availability

Custom scripts producing the analyses and figures will be made available in an OSF repository upon publication. EEG data are available in an OSF repository.

Readers seeking access to the stimulus set should contact Anat Perry from The Hebrew University of Jerusalem. Access will be granted to named individuals in accordance with ethical procedures governing the reuse of sensitive data.

## Acknowledgements

We would like to thank Shira Inbar for illustrating the EEG cap, Shlomi Fridge for partaking in the linguistic annotation effort, Meir Horovitz for partaking in the EEG preprocessing effort, Nir Ofir for developing LIMO-related scripts and plotting scripts, and providing comments on the manuscript. For discussions that improved this study we thank Nadav Matalon and members of the Brain, Attention and Time lab.

## Supplementary Information

**Figure S1.**
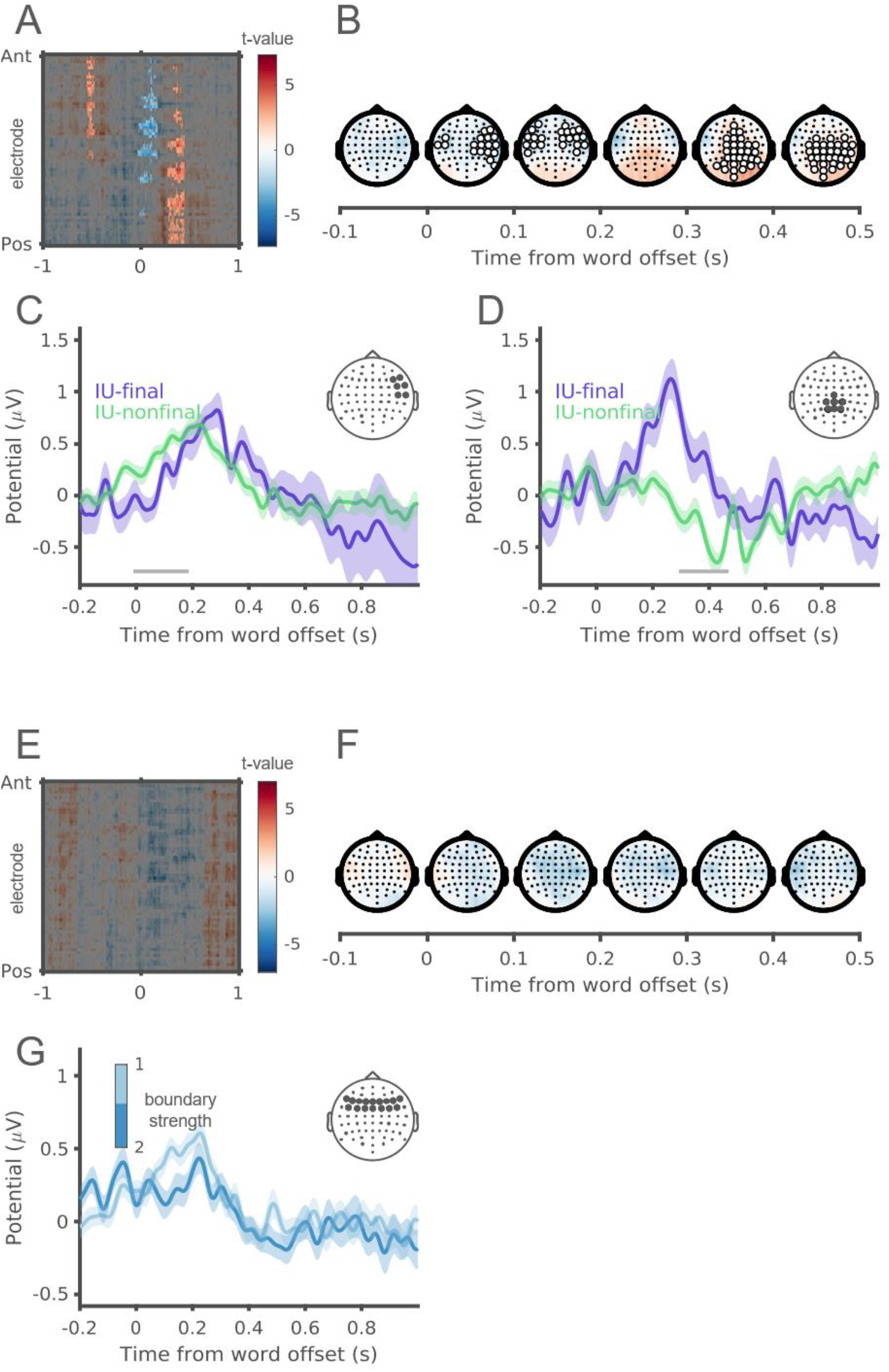
EEG amplitude depends on IU closure and boundary strength (non-standardized variables). (A) The full spatiotemporal results for the IU closure effect at zero boundary strength. A t-value is calculated for the observed difference between IU-final and IU-nonfinal EEG responses at each electrode and timepoint in a 2-second window around word offset. Cluster-based bootstrap tests reveal significant clusters at the group level (unmasked). (B) Topographical distributions of the IU-closure contrast t-values, from -100 ms to 500 ms around word offset in 100 ms steps. Electrodes in significant clusters are highlighted in white if they span 15 consecutive milliseconds. For words with zero boundary strength, words that close an IU are associated with a centroparietal positivity preceded by a right-to-bilateral anterior negativity compared with words that do not close an IU. This was supported by the significant effect of IU closure (Figures 3A-D; negative cluster extending from -12 to 188 ms following word offset, p = 0.003; positive cluster extending from 291 to 467 ms following word offset, p < 0.001). There was an additional positive cluster preceding the negativity, extending from -537 until -453 ms relative to word offset, p = 0.02). (C) ERP traces in response to IU-final and IU-nonfinal words with zero boundary strength score (purple and green, respectively; as opposed to mean boundary strength scores, in Fig 3C), illustrating the right-anterior-negative cluster. The traces show the grand average over the EEG electrodes highlighted in the inset topography. Shaded ribbons correspond to ±1 SEM. The horizontal gray bar marks the timepoints over which this cluster was significant in the GLM model. (D) ERP traces in response to IU-final and IU-nonfinal words with zero boundary strength score (purple and green, respectively), illustrating the centroparietal positive cluster. The traces show the grand average over the EEG electrodes highlighted in the inset topography. Shaded ribbons correspond to ±1 SEM. The horizontal gray bar marks the timepoints over which this cluster was significant in the GLM model. (E) The full spatiotemporal results for the effect of boundary strength within IU-nonfinal words. A t-value is calculated for the estimated change in EEG response against zero at each electrode and timepoint in a 2-second window around word offset. Cluster-based bootstrap tests reveal no significant effect of boundary strength within IU-nonfinal words, therefore no clusters are unmasked. (F) Topographical distributions of the effect of boundary strength within IU-nonfinal words, from -100 ms to 500 ms around word offset in 100 ms steps. There was no significant effect of boundary strength within IU-nonfinal words, therefore no electrodes are highlighted. (G) ERP traces in response to IU-nonfinal words with different levels of boundary strength, illustrating a trend towards an anterior negative cluster. In light of the lower variability in boundary strength for IU-nonfinal words (see Figure 2A), only two different levels are presented, roughly corresponding to a median split of boundary strength scores within IU-nonfinal words. The traces show the grand average over the EEG electrodes highlighted in the inset topography. Shaded ribbons correspond to ±1 SEM.

### Appendix S1: Example transcript

In the following we present the transcript of one of the stories that served as stimuli in the EEG experiment. We provide a broad phonetic transcription of the Hebrew text, and note the following deviations from the standard IPA conventions:

`S` – substitutes /ʃ/

`x` – substitutes /χ/

`c` – substitutes /ʦ/

`‘` – substitutes /ʔ/, a glottal stop which is often elided

We follow the orthographic conventions in Hebrew for defining words in the transcription. We use hyphens to indicate cases in which orthographic conventions treat separable morphemes as single words (e.g., `ha-miSpaxa` ‘the family’). Comments in double brackets, `((xxxx))`, describe nonverbal material. Occasional curly brackets, `{00:30:00}`, indicate recording time. The prosodic-based segmentation into Intonation Units (IUs) is represented by a break in line, such that each numbered line corresponds to a single IU. We provide a literal English translation to each IU.

`Stimulus 03:`

1. `ehh` uhh
2. `hayti xayal` I was a soldier
3. `be` in
4. `ba-cava` in the army
5. `ba-tironut` in training
6. `((breath))`
7. `ve` and
8. `hayta li bat doda` I had a cousin
9. `Se` that
10. `lo hirgiSa tov` didn’t feel well
11. `((breath))`
12. `hamon hamon zman` for a very very very long time
13. `((breath))`
14. `ve` and
15. *be-’eizeSehu* Slav at some point
16. `ha-macav Sela` her situation
17. `hitxil lehitdarder` started deteriorating
18. `ze kvar ehh` it already uhh
19. `hitxil laruc` started running
20. `((breath))`
21. `bein ha` between the
22. `bein ha` between the
23. `bnei dodim` cousins
24. `ha-miSpaxa` the family
25. `Se` that
26. `ha-macav mitdarder` the situation is deteriorating
27. `((breath))`
28. `ve` and
29. `‘ani zoxer Se-lo hayti mamaS me’orav` I remember that I wasn’t really involved
30. `be-kol ha` in the whole `{00:30:00}`
31. `emm` umm
32. `be-kol ha-tahalix` in the whole process
33. `ve-ba-hitdarderut Sela` and in her deterioration
34. `lo hayti me-’ele Se-mevakrim ‘ota yoter midai` I wasn’t one of those who visited her so much
35. `((breath))`
36. `‘az lo yadati ‘ex ‘ani ‘agiv kSe` so I didn’t know how I would react when
37. `ehh` uhh
38. `hi hayta xola be-maxala sofanit` she was terminally ill
39. `‘az lo` so I didn’t
40. `lo yadati ‘ex ‘ani ‘agiv` I didn’t know how I would react
41. `kSe-ze ykre` when it would happen
42. `((breath))`
43. `‘az ze` so it
44. `hiftia ‘oti` surprised me
45. `((breath))`
46. `ve` and
47. `be-’eizeSehu Slav ba-tironut` at some point in training
48. `emm` umm
49. `kibalti sms` I got a text message
50. `Se-ha-macav mamaS hitdarder` that the situation really deteriorated
51. `tevakSu bakaSot yeci’a` ask for exit leaves
52. `ke’ilu` like
53. `tavo’u levaker` come visit
54. `lehagid ehh` to say uhh
55. `Salom` goodbye
56. `bet xolim ve-ze` hospital etc.
57. `((breath))`
58. `‘az ehh` so uhh
59. `‘ani zoxer Se-’ani yaradeti la` I remember that I went down to the
60. `mefaked` commander
61. `levakeS ehh` to ask uhh `{01:00:00}`
62. `‘et-bakaSat ha-yeci’a` for the exit leave
63. `hisbarti lo ma kara` I explained to him what happened
64. `((breath))`
65. `emm` umm
66. `‘az hu ‘amar li Se-hu yivdok ‘et-ze` so he told me that he would look into it
67. `‘az ‘aliti be-xazara lemala la-pelefon` so I went back up to the cellphone
68. `kedei ehh` to uhh
69. `laxzor` go back
70. `hayti be-xol ‘ofen be-S’at taS` I was anyway in the after hours
71. `((breath))`
72. `laxzor lir’ot ma ma ha-macav` to go back and see what what’s the situation
73. `((breath))`
74. `emm` umm
75. `ve-’az hodi’u Se-hi kvar niftera` and then I was told that she already passed away
76. `ve-’az ehh` and then uhh
77. `ha-tguva Seli ‘ani zoxer hayta` my response I remember was
78. `((breath))`
79. `ehh` uhh
80. `Se` that
81. `paSut` just
82. `nafal li ha-pelefon me-ha-yad` the cellphone fell from my hand
83. `((breath))`
84. `ve` and
85. `paSut` just
86. `hirgaSti` I felt
87. `mamaS mamaS mamaS nora` really really really bad
88. `ve` and
89. `nirdema li yad smol` my left hand became numb `{01:30:00}`
90. `lo hayta li txuSa be-yad smol` I had no sensation in my left hand
91. `((breath))`
92. `hitxalti livkot` I started crying
93. `ve-lo hiclaxti lehiStalet ‘al ‘acmi` and didn’t manage to control myself
94. `((breath))`
95. `‘ani zoxer Se-haynu` I remember that we were
96. `‘axra’im` responsible
97. ((click)) *‘al ha-neSakim Selanu* ((click)) for our rifles
98. `ki haynu tironim` because we were trainees
99. `haynu crixim lakaxat ‘otam le-kol makom` we had to take them everywhere
100. `((breath))`
101. `ve-hayti kol kax` and I was so
102. `emm` umm
103. `Sabur` broken
104. `Se-Saxaxti ‘et-ha-neSek ba` that I forgot the rifle in the
105. `mita` bed
106. `((breath))`
107. `‘az ehh` so uhh
108. `hayta xavaya mamaS lo pSuta` it was really not an easy experience
109. `((breath))`
110. `ve` and
111. `emm` umm
112. `zehu` that’s it
113. `‘az ha-mefaked` so the commander
114. `lamrot kol ha` despite all the
115. `distans` distance
116. `Se-haya be-’oto zman` that was at the time
117. `lakax ‘oti le-sixa` took me for a conversation
118. `((breath))`
119. `ve` and
120. `diber ‘iti` spoke with me
121. `ve` and
122. `nisa le’at le’at lehargia ’oti` tried slowly to calm me down
123. `((breath)`
124. `‘aval xad maSma’it ehh` but definitely uhh `{02:00:00}`
125. `‘exad ha` one of the
126. `xavayot ha-kaSot` difficult experiences
127. `((breath))`
128. `ehh` uhh
129. `‘az zehu` so that’s it

